# Know Today, Know Tomorrow: Ensemble Forecasting of Wildlife Sightings from Temporal Dynamics

**DOI:** 10.64898/2025.12.09.693309

**Authors:** Takeshi Honda, Chinatsu Kozakai

## Abstract

1. Forecasting encounters between humans and large carnivores has largely relied on mechanistic models driven by causal factors such as food resources and weather. However, for short-term forecasting these approaches implicitly require unrealistically detailed real-time data on many covariates and an almost complete understanding of the underlying causal pathways. As a result, they offer little practical support for short-term, operational decision-making.
2. We developed a short-term forecasting system that predicts end-of-month cumulative bear sightings from the beginning of each month by exploiting temporal autocorrelation without mechanistic assumptions, using an ensemble of multiple components: (i) sequential estimation of daily sighting rates via a non-stationary Poisson process, (ii) seasonal baselines with ratio-based corrections from previous months, and (iii) rule-based transitions among components as daily sightings accumulate.
3. Applied to Asiatic black bear (Ursus thibetanus) sighting records from two Japanese regions differing 18-fold in encounter frequency (maximum monthly counts: 83 vs. 1490) and with contrasting seasonal peaks, the ensemble achieved correlations of ≥0.8 between predicted and observed month-end totals from day 1, increasing to ≥0.98 by day 20 and substantially outperforming a null model that assumed no seasonal or interannual variation (ΔAIC: 477−652).
4. After controlling for baseline spatial risk and for the region-wide daily bear-forecast level (temporal risk) provided by our ensemble, we detected strongly localized short-term recurrence in bear encounters: prior sightings increased encounter probability within 500 m for up to 3 days, with rapid decay in space and time.
5. *Synthesis and applications*. This observation-based ensemble demonstrates that temporal dynamics alone can approach the practical limits of short-term predictability of wildlife encounter rates, without relying on detailed environmental covariates or extensive new data collection. By quantifying both when (daily risk levels) and where (localized hotspots around recent sightings) encounters are most likely, the system offers wildlife agencies and residents an immediately implementable tool for issuing targeted warnings, adjusting outdoor activities, and reducing human injuries in regions experiencing increasing human–carnivore conflict.

## 1 Introduction

Forecasts shape our world (Dietze et al., 2018; Olsson et al., 2025). Weather forecasts allow us to decide whether to carry an umbrella, while predictions of declining fish stocks inform limits on harvest (Okamura et al., 2018; Tsikliras & Froese, 2019). Such forward-looking decisions rely fundamentally on forecasting. Ecology is no exception: predictions of increasing large ungulate populations and the accompanying rise in human-wildlife conflict guide decisions to control population size (Conover, 2001; Reidinger & Miller, 2013). For large carnivores such as Asiatic black bears *Ursus thibetanus*, forecasting is closely linked to human safety. Recent incidents have involved fatal and severely injurious attacks on people engaged in routine activities such as commuting, farming, walking, or shopping, underscoring how easily ordinary daily activities can be disrupted (Akita Prefecture, 2025b; McCurry & Corlett, 2025). Severe bear attacks have led authorities in some regions to ask residents to limit nonessential outdoor activities when bears are repeatedly sighted near residential areas.

Given the central role played by forecasts, it is worth asking what conceptual and computational frameworks have been used to produce them. The fundamental approach is to explicitly model the mechanisms generating phenomena—cause and effect (Oidtman et al., 2021; Yates et al., 2018). Weather forecasts model atmospheric thermodynamics, while infectious disease predictions incorporate human movement and infection rates (Kraemer et al., 2020; Krishnamurthy, 2019). Wildlife forecasts follow the same principle, building models that assume causal relationships (Burns et al., 2003; Honda & Kozakai, 2020; Molnár et al., 2011). In predicting injuries from bear encounters, for example, models treat food scarcity—mast crop failures, pine seeds, or salmon shortages—as causal drivers (Artelle et al., 2016; Mattson et al., 1992; Oka et al., 2004). These mechanistic models have attracted extensive research, as they provide frameworks not only for prediction but also for understanding ecosystems (Beckage et al., 2011; Yates et al., 2018).

However, short-term predictive accuracy remains challenging for mechanistic models. A review of population models revealed that many sophisticated models failed to outperform a simple null model—the assumption that “next year = this year” (Ward et al., 2014). In COVID-19 mortality forecasts, approximately 30% of complex models performed worse than this null model (Cramer et al., 2022). As George Box famously noted, all models are wrong—they can never represent the true system. Yet, even when we attempt to approximate the truth by adding more detail or complexity, predictive accuracy does not necessarily improve (Burnham & Anderson, 2004; Makridakis & Hibon, 2000). In wildlife systems, ecological processes are only partially observed, making it difficult to construct complex mechanistic models. Furthermore, the available evidence is often insufficient to determine which drivers are most influential, limiting the development of even modestly simplified models. Bear sightings provide a clear illustration of this structural limitation. Models based on autumn food availability (e.g., hard mast, pine seeds, salmon) fail to explain sighting variations earlier in the year. Models incorporating spring frost effects (Honda, 2013) account for impacts on flowering and fruit production but ignore insect prey. Even when combining autumn food and frost effects, models fail during extreme drought years if they neglect summer heat stress on vegetation. No model can ever be complete (Burnham & Anderson, 2004; Green & Armstrong, 2015). And in systems like bear sightings—where key drivers cannot be clearly identified—there is no reliable basis for selecting which factors to retain, making meaningful simplification impossible.

An alternative exists. Short-term forecasting enables a different approach that requires no mechanistic assumptions (Pennekamp et al., 2019; Petchey et al., 2015). Bear sightings exhibit temporal continuity: this week’s rate predicts next week’s. Simple moving-average methods utilize this continuity but are inherently retrospective. A 10-day moving average, for instance, effectively estimates the situation as it was approximately 5 days ago, not as it is today. While such smoothing is acceptable for describing historical trends, it is inadequate for forecasting rapidly shifting risks. This immediacy is particularly critical for predicting human-bear conflicts, which are characterized by intense seasonal fluctuations and sudden surges. Consequently, effective wildlife forecasting requires “nowcasting”—estimating the true underlying risk at the present moment. The non-stationary Poisson process offers a promising framework for this purpose. By quantifying how the sighting intensity changes through time, this approach represents encounter risk as a smoothly varying state—a structure that allows increases or decreases to be detected instantaneously, rather than only after past values have been averaged. Without such forecasts, residents cannot objectively judge how dangerous it is to go outside today. A reliable daily forecast that quantifies current bear encounter risk would enable decision-making similar to that based on weather forecasts. Residents could assess whether today’s risk justifies limiting outdoor activities, and to what degree. This is what responsive short-term forecasting must provide.

While the non-stationary Poisson process is a powerful tool, relying exclusively on a single non-mechanistic model is akin to characterizing wildlife dynamics using only one predictor. Time-series data offer rich possibilities for representing temporal structure—seasonal cycles, interannual fluctuations, and short-term trends can each yield complementary insights. Consequently, integrating diverse non-mechanistic models that capture distinct facets of these temporal dynamics promises greater efficacy. In recent years, information-theoretic approaches such as AIC have highlighted that multiple competing models can simultaneously garner strong empirical support (e.g., models with ΔAIC < 2). This recognition—that multiple models can be concurrently valid—aligns naturally with ensemble forecasting, in which combining diverse models offsets individual biases and enhances predictive accuracy (Bates & Granger, 1969; Leutbecher & Palmer, 2008). Although ensemble forecasting is standard practice in meteorology and epidemiology, its potential regarding terrestrial mammals remains largely unexplored (Molnár et al., 2010). Our study represents a first step toward realizing this potential by constructing an ensemble of non-mechanistic bear activity forecasts, with the non-stationary Poisson process serving as a critical component for capturing day-to-day fluctuations in sighting rates.

## 2 Methods

Our forecasting approach uses an ensemble of several predictive methods (Fig. 1; details follow). Here, an ensemble means combining forecasts that rely on different assumptions or information sources so that the strengths of each method complement one another. Predictions based on a single model depend heavily on that model’s assumptions and error structure, and they can fail when conditions differ from what the model expects. By contrast, an ensemble brings together multiple forecasts and averages out their individual biases, providing a more reliable overall prediction. In the following sections, we explain each step of the ensemble in the order shown in Fig. 1. All statistical analyses were conducted in R version 4.5.1 (R Core Team, 2025). We fitted Bayesian models in Stan via the R package rstan (Stan Development Team, 2024).”

**Fig. 1.**
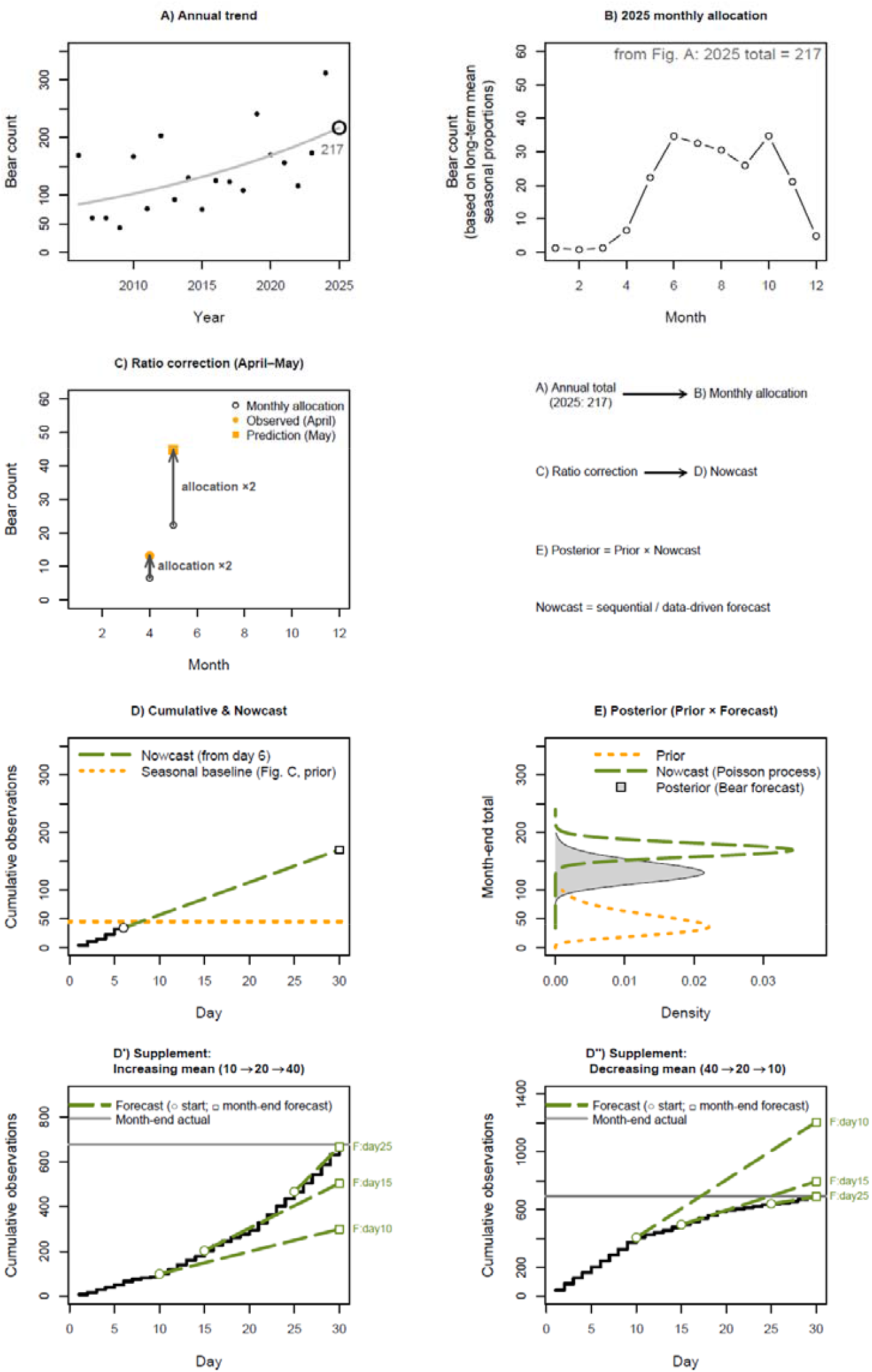
Structure of the ensemble forecasting framework for daily bear sightings. (A) Long-term trend of annual sightings (GLMM with a common slope and random intercepts across prefectures). (B) Monthly allocation for 2025 derived from the annual total in (A). (C) Adaptive correction that adjusts monthly allocation based on the observed-to-predicted ratio from the previous month (April → May example). (D) Daily cumulative forecast based on a non-stationary Poisson process (nowcast from day 6). (E) Posterior forecast (bear forecast) obtained by Bayesian ensemble of the prior (weighted prediction from monthly allocation and ratio correction, Figs. 1B-C) and the nowcast (sequential Poisson process, Fig. 1D). (D′, D″) Illustrations of non-stationary Poisson dynamics with increasing (D′) and decreasing (D″) daily means, respectively, showing forecast trajectories from days 10, 15, and 25.

### 2.1 Data

We used sighting reports of Asiatic black bears from two regions of Japan: a central region (Yamanashi Prefecture) and a northern region (Akita Prefecture). Both datasets are publicly available and can be downloaded from each prefecture’s website (Akita Prefecture, 2025a; Yamanashi Prefecture, 2025). In both focal regions, residents report bear sightings to their municipal offices and/or the police. These reports are not casual observations; they are requests for assistance, because a nearby bear poses a clear safety concern. In the northern region, for example, 34 people were injured by bears during the peak sighting period—a single month—numbers that are substantial even when compared with traffic-related injuries (122 cases) during the same period. As a result, the reporting rate is very high, and the records compiled by the prefectural governments provide reliable information on when and where bears were seen. Prefectural governments organize these reports and record the date, location, and circumstances of each sighting. All components of the forecasting framework in Fig. 1 were based on these records. The available data spanned 2019–2024 in the central region and 2022–2024 in the northern region. Sightings differed greatly between regions: the maximum monthly count reached 83 in the central region but 1490 in the northern region (≈18-fold difference). Because the two regions vary widely in geography and sighting intensity, consistent forecasting performance across both provides a strong indication of model robustness.

To estimate long-term trends, we also used monthly sighting data from 30 prefectural regions across Japan. The length of the time series differed by prefecture, covering 1997–2024. These nationwide data were used to estimate the long-term trend in Fig. 1A and the seasonal pattern in Fig. 1B. Because bears do not inhabit all prefectures, the number of regions included in the analysis is smaller than the total number of prefectures in Japan.

### 2.2 Baseline for forecast (Fig. 1A–B)

To construct a forecasting baseline, we first described the long-term national pattern in annual bear sightings. This baseline is meant only to set the overall level of expected sightings; it is not designed to follow short-term ups and downs. Because annual sighting counts can shift gradually over time, the trend needs to be smooth enough to avoid reacting to year-to-year noise while still capturing the general direction of change. For this reason, we used a simple linear form rather than nonlinear or periodic alternatives. We fitted a generalized linear mixed model (GLMM; Poisson family) with year as a fixed effect and prefecture identity as a random intercept. The slope of the year effect was shared across prefectures, while differences in average sighting levels were absorbed by the random effect. This structure provides a stable nationwide trend that serves only as a starting point; more detailed, region-specific adjustments are added in later steps (Fig. 1B–E). The model used annual sighting counts for each prefecture, noting that the available number of years differed among prefectures. The analysis was implemented in R using the glmmML package, which provided reliable convergence for this simple mixed-effects specification. The resulting baseline prediction for the central region in 2025—the value used in Fig. 1A—was 217.

Next, we allocated the baseline annual value (217) to individual months using the nationwide seasonal pattern of bear sightings (Fig. 1B). To obtain this pattern, we fitted a separate GLMM to nationwide monthly sighting counts, including month as a fixed effect and prefecture identity as a random intercept. The month effect yields a set of relative monthly ratios that capture the typical seasonal variation in sighting activity. Applying these ratios to the annual baseline produced monthly values that represent expected sighting levels under ordinary conditions. We refer to this baseline as F_base, which serves as the starting point that is later refined by the ratio correction and the short-term Poisson process (Fig. 1C–E).

### 2.3 Ratio correction (Fig. 1C)

The second component of the forecast is an empirical ratio correction, denoted F_ratio, which carries over the forecasting error from the previous month (Fig. 1C). The correction is defined as the ratio of observed to predicted sightings in the previous month:

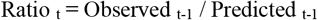

This ratio is then applied to the baseline forecast for the current month. For example, if the previous month’s ratio was 2, we assume that a similar degree of underprediction will continue, and the current baseline value (F_base) is multiplied by 2. This approach is intentionally simple. It is not a mechanistic model of how errors accumulate, but an empirical way to reflect the common tendency for forecasting errors to persist from one month to the next. In doing so, it approximates the effect of temporal autocorrelation in sighting intensity and allows recent deviations to inform expectations for the coming month. Such simplicity is useful here, because ensemble forecasting can benefit from combining several clear, easy-to-understand components, each capturing a different part of the overall pattern, rather than relying on a single, highly complex model.

### 2.4 Poisson process (Fig. 1D)

The third component uses a non-stationary Poisson process to capture day-to-day changes in sighting intensity (Fig. 1D). This approach treats daily sighting counts as Poisson-distributed random variables whose mean rate varies smoothly through time. The rate—representing expected sightings per day—is modeled as a random walk on the log scale: each day’s rate equals the previous day’s rate plus a small random increment. This structure allows the model to track increasing or decreasing trends in sighting activity as they develop, rather than averaging over past observations. The model forecasts cumulative monthly counts, which by definition can only increase: once a bear is sighted, that observation remains in the record. However, the *rate* at which sightings accumulate can change. When sighting intensity rises during a month (Fig. 1D′), forecasts issued early in the month underestimate the final total. When intensity declines (Fig. 1D″), early forecasts overestimate. As the month progresses and more observations become available, forecasts converge toward the actual trajectory.

The model was implemented in Stan (Appendix Code 1). We denote this forecast component as F_PoissonP.

### 2.5 Ensemble: F_base and F_ratio (Fig. 1E)

The next step combines F_base (Fig. 1B) and F_ratio (Fig. 1C) into a single weighted prediction. This weighting step is not shown explicitly in Fig. 1 for simplicity. The ratio correction can become unstable when predicted counts are small. In months with few sightings, the denominator in the ratio approaches zero, causing erratic behavior. To avoid this, we weighted F_ratio according to the previous month’s predicted count:

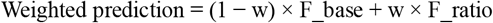

where the weight w is:

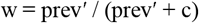

and prev′ = log(previous month’s prediction + 1). The log transformation with +1 prevents numerical issues when predictions are zero. The constant c controls how the weight transitions between F_base and F_ratio. We estimated c by minimizing the squared prediction error across all prefectures and months using nonlinear least squares (optim in R). When predicted counts are large, the ratio correction is reliable and w approaches 1, giving more weight to F_ratio. When counts are small, w approaches 0 and F_base dominates. The parameter c determines the smoothness of this transition. In Fig. 1E, the “prior” represents the distribution of this weighted prediction—the expected distribution of month-end counts before incorporating daily observations. We modeled this as a gamma distribution with mean equal to the weighted prediction and variance estimated from the correlation between observed and predicted monthly counts. Specifically, we defined variance as a decreasing function v(r) of the correlation coefficient r, where v(r) = 0 when r = 1 and increases as r decreases. We chose the gamma distribution because expected counts are positive and continuous.

### 2.6 Ensemble blending rule (Fig. 1E)

The final forecast blends the weighted prediction with F_PoissonP. Early in the month, the forecast relies on the weighted prediction; later, it shifts toward F_PoissonP as more observations accumulate. The weighted prediction operates at a monthly scale, capturing seasonal patterns adjusted for recent trends, while F_PoissonP operates daily, tracking fluctuations in the current sighting rate.

Early in the month (days 1–10), few observations are available and the Poisson process estimate is unstable, so the forecast relies entirely on the weighted prediction. Through mid-month (days 11–19), we transition linearly:

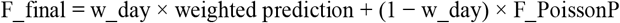

where w_day decreases from 0.9 on day 11 to 0.1 on day 19. From day 20 onward, the forecast uses only F_PoissonP, projecting the current rate through month-end.

We include one exception for months that develop faster than expected. If cumulative observed counts reach 70% of the weighted prediction before day 20, the forecast switches immediately to F_PoissonP. This early-switch rule recognizes that the month is progressing at an unusually high rate, and continuing to weight the seasonal baseline would underestimate the month-end total. The 70% threshold aligns approximately with two-thirds of the month while providing a margin against premature switching. This rule-based approach ensures that the forecast remains interpretable. The transition logic is transparent, the same rules apply across all regions, and the system avoids region-specific parameter tuning. Performance across both study regions is evaluated in Results.

### 2.7 Spatial extension: short-term risk around previous sightings

Bear sightings sometimes cluster in space and time. When a bear is sighted at one location, subsequent sightings often occur nearby within days. This clustering allows us to distinguish between two different processes. First, baseline spatiotemporal risk. Some locations are inherently risky (e.g., near forest cover), and some periods carry elevated activity across the region (captured by F_final). Sightings may cluster in these contexts simply because favorable habitat and seasonal conditions attract bears generally, regardless of specific recent events. Second, short-term repeat sightings. A sighting on Monday at one location, followed by another sighting 500 meters away on Tuesday, suggests mechanisms beyond baseline risk. Because bears have finite movement rates, an individual sighted at one location may remain in or return to nearby areas over subsequent days. This produces spatiotemporal dependence that cannot be explained by baseline risk alone—it depends on the specific history of recent sightings.

We tested for short-term clustering using a logistic regression model that controls for baseline risk. The response variable was bear presence (1) or absence (0). Presence locations were the observed sightings; absence locations were pseudo-absence points randomly generated across the entire study region and study period. This design allows us to ask a simple question. After accounting for where and when bear sightings are generally likely to occur (baseline risk), are locations closer to a recent sighting still more likely to host a bear? The model included four components. First, spatial risk: the baseline probability that a location attracts bears, estimated from distances to forest, roads, and settlements, with a spatially autocorrelated Gaussian random field to capture residual spatial structure (glmmfields package in R). Second, temporal risk: the daily risk level from our forecasting model (F_final), reflecting region-wide activity on that date. Third, lag: the number of days since the nearest previous sighting (evaluated from 0 to 7 days). Fourth, distance: the distance to that nearest previous sighting. We also included an interaction term (lag × distance) to test whether the clustering effect decays with both time and space. For interpretation, we expressed model predictions as relative occurrence risk compared with a reference condition of being 10 m from a same-day sighting (lag = 0, distance = 10 m). Essentially, this approach looks at what remains unexplained. By explicitly modeling why a location is typically dangerous (spatial baseline) and why a day is typically active (temporal baseline), we filter out the usual environmental risks. If being close to yesterday’s sighting still predicts today’s danger even after this rigorous check, it implies that the event itself is creating a temporary “hotspot”—a localized spike in risk driven by what just happened, rather than by the environment alone. The key question is whether lag and distance predict presence after accounting for spatial and temporal baselines. A significant effect of proximity—short lag and small distance—indicates that recent sightings elevate risk locally, beyond what baseline conditions would predict.

## 3 Results

### 3.1 Forecasting accuracy

We achieved high predictive accuracy for bear sightings. An overview of forecasting performance (Fig. 2) shows that accuracy improved progressively from early to late in each month. Throughout the season, our forecasts consistently outperformed a simple intercept-only null model with no seasonal or interannual effects (ΔAIC on day 1: 477.2, day 10: 489.7, day 20: 651.8). Our ensemble forecast operated as follows. Early in each month, predictions relied on (i) seasonal patterns based on historical monthly averages (Fig. 1B) and (ii) a ratio correction that carried forward the previous month’s forecasting error (Fig. 1C). As daily observations accumulated, the forecast transitioned progressively toward (iii) the non-stationary Poisson process (Fig. 1D). Component weights were predetermined by rules reflecting information availability over time, not optimized post hoc from observed data. Under this structure, forecasting accuracy in the central region (Yamanashi Prefecture) exceeded a correlation coefficient of 0.98 after day 21, when the system fully transitioned to the Poisson process. In the northern region (Akita Prefecture), correlations reached 0.98 by day 17. High accuracy was thus achieved regardless of location or sighting intensity.

**Fig. 2.**
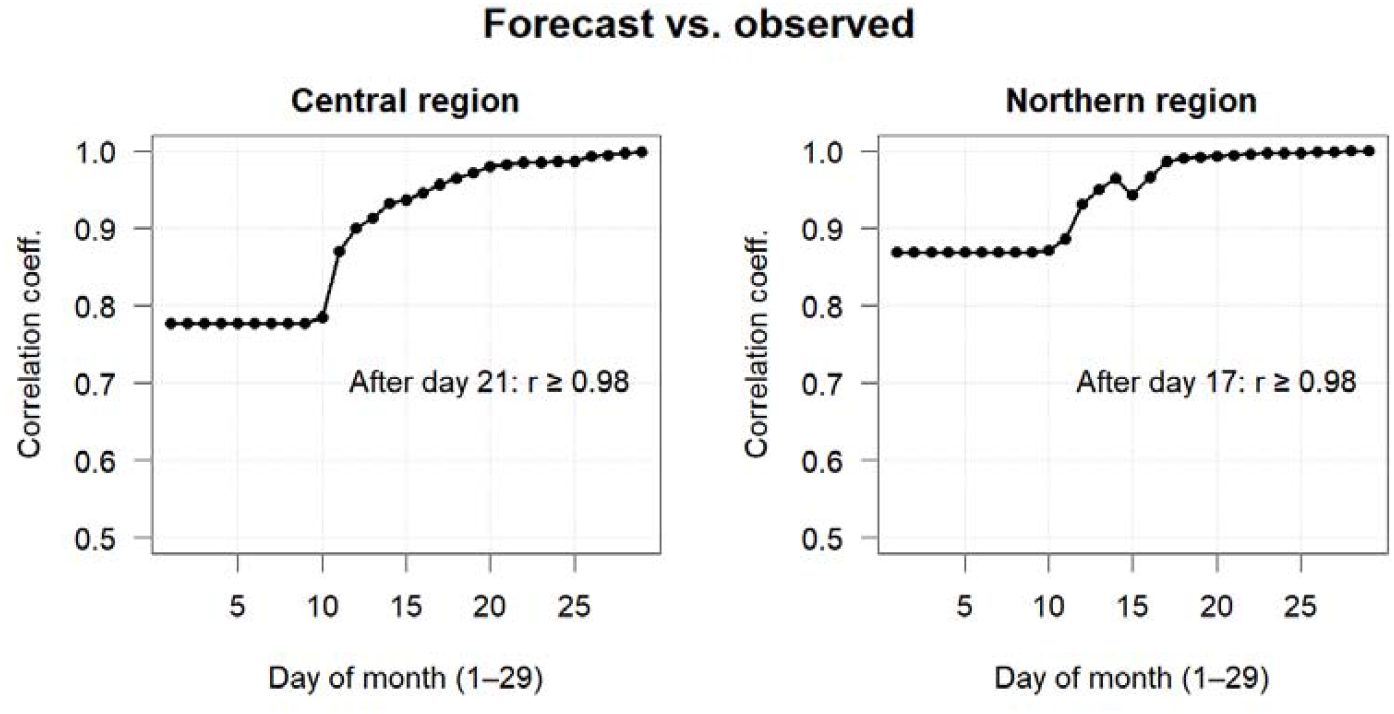
Correlation between forecasted and observed monthly bear sightings as a function of day within the month. Left: central region (Yamanashi Prefecture). Right: northern region (Akita Prefecture). Distinct phases are visible as the system shifts from initial estimates excluding the Poisson process to updated forecasts incorporating the Poisson process, demonstrating region-specific timelines for achieving reliable accuracy.

Examining forecasts across individual months (Fig. 3), predictions tracked observed month-end totals closely in both high-activity and low-activity years across both regions. The bear index—monthly sighting counts standardized to each region’s maximum value during the study period (83 for central, 1490 for northern region)—illustrates forecast trajectories throughout the year. In years with lower activity (2022 in both regions), forecasts converged rapidly to final values from early in each month. In high-activity years (2024 for central, 2023 for northern region), early-month predictions showed somewhat larger deviations during peak months, but by mid-month the Poisson process adapted to the current sighting rate and forecasts remained accurate even during the most intense outbreak periods (e.g., June in Yamanashi 2024, October in Akita 2023). Notably, although the seasonal baseline was estimated as a single nationwide pattern, the ensemble still captured different peak seasons in each region—early summer in the central region and autumn in the northern region. This suggests that region-specific dynamics were captured by the ratio-correction and Poisson process components, despite the use of a common seasonal baseline. Regional differences in early-month accuracy reflected differences in sighting counts (Appendix Fig. 1, P < 0.001, GAM, quasi-binomial): regions with higher historical sighting numbers showed higher early-month accuracy. Our ensemble assigns greater weight to the ratio correction when sighting numbers are large, following the statistical principle that estimation reliability improves with data abundance.

**Fig. 3.**
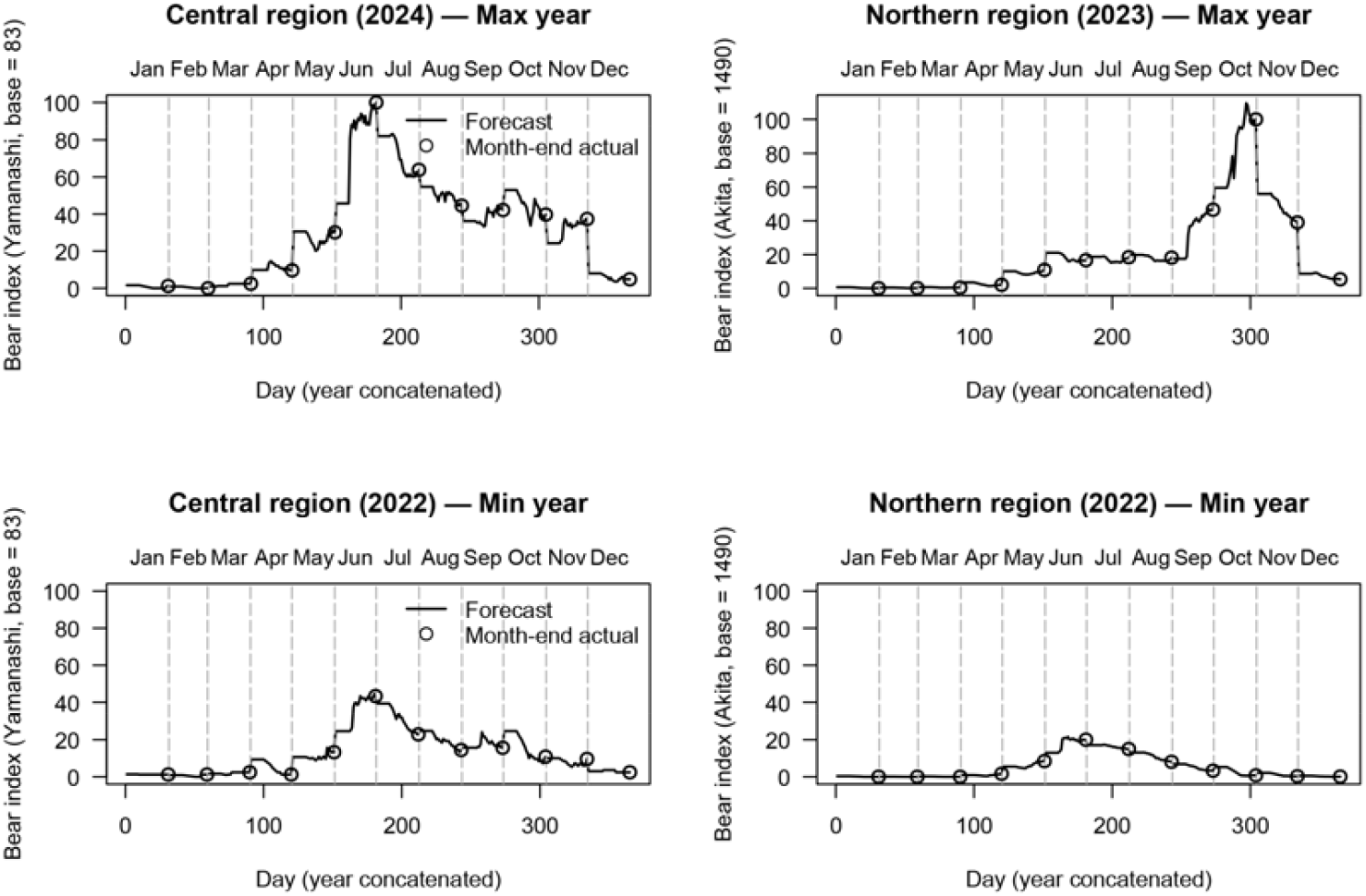
Daily forecast progression across an entire year in the central region (Yamanashi Prefecture, left panels) and northern region (Akita Prefecture, right panels). Lines show forecasted cumulative monthly totals updated daily; circles show month-end observed values. These plots substantiate the high accuracy shown in Fig. 2 by detailing actual forecast behavior. Early-month predictions (lines) often closely approximate final outcomes (circles), and any initial deviations rapidly diminish. Notably, by mid-month, forecasts align almost perfectly with observed totals, even during extreme outbreak years (upper panels). The bear index standardizes monthly sighting counts to each region’s maximum during the study period (Yamanashi: 83 in June 2024; Akita: 1490 in October 2023). Upper panels show maximum-activity years; lower panels show minimum-activity years. The Poisson process enables forecasts to adapt within each month as sighting rates shift, ensuring robust performance across diverse seasonal patterns and outbreak scales.

### 3.2 Short-term inducement effect of prior bear sightings

We quantified the short-term influence of prior sightings using logistic regression, after controlling for baseline spatial risk (Appendix Fig. 2) and temporal risk (Fig. 3). The analysis compared observed sighting locations (presence) against pseudo-absence points randomly generated. Predictors included (1) baseline spatial risk, (2) baseline temporal risk, (3) time lag (days since the nearest prior sighting), and (4) distance to that prior sighting.

The results revealed a distinct and statistically significant clustering effect (P < 0.001). However, this elevated risk was highly localized in both space and time (Fig. 4). Relative to the risk at the immediate location of a sighting on the same day, the relative probability of a subsequent encounter dropped to approximately 23% at a distance of just 100 m. The effect also decayed rapidly over time; even at the exact same location, the elevated risk diminished to approximately 4% after 7 days. These findings indicate that while sightings do cluster, the window of heightened local risk is extremely narrow.

**Fig. 4.**
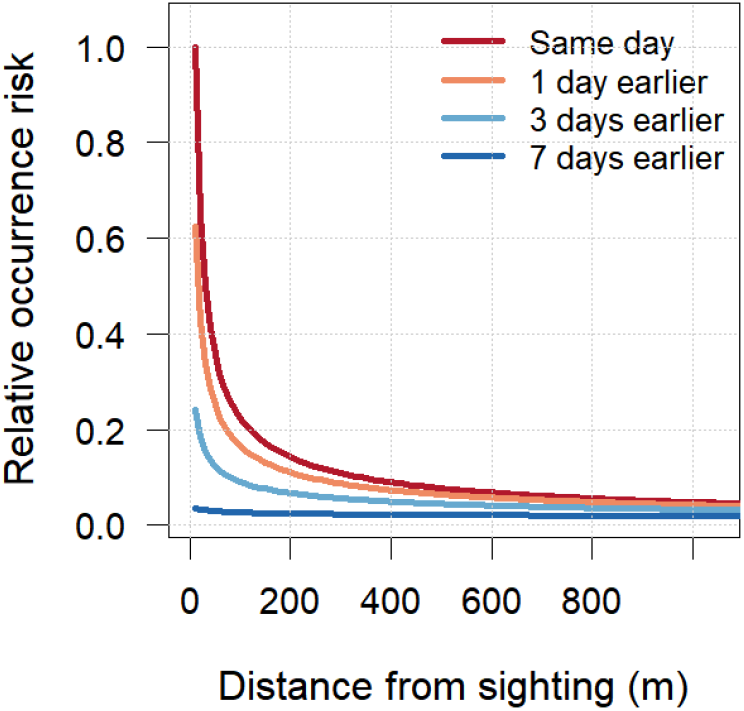
Impact of consecutive bear sightings. This figure illustrates the relative risk of one sighting triggering another, estimated after controlling for baseline geographic (Appendix Fig. 2) and temporal (Fig. 3) risks. This step is crucial to distinguish the specific impact of a recent sighting from generally high-risk areas or periods. We normalized the risk to 1.0 for a recurrence at the exact same location and day. By isolating the inducement effect of the event itself, these curves show, for example, the level of risk remaining the next day at various distances from the initial sighting.

## 4 Discussion

### 4.1 Accurate forecasts without mechanistic assumptions

Our ensemble forecasts demonstrated high accuracy. This success stems from complementarity across time: each model excels under different data conditions. Early in the month, when current observations are scarce and the Poisson process remains unstable, seasonal patterns and ratio-based corrections provide reliable forecasts. Later, as sightings accumulate, the Poisson process captures the ongoing rate of encounters, projecting it forward to predict monthly totals. This combination substantially outperformed the null model (ΔAIC: 477–652), addressing the persistent problem that many ecological forecasts fail to beat simple baselines (Cramer et al., 2022; Ward et al., 2014).

Monthly forecasts yielded correlations exceeding 0.8 during the first ten days, rising to approximately 0.98 from mid-month onward. Even years with mass sightings showed close agreement with observations by mid-month. Comparable predictive performance was achieved across regions with markedly different sighting magnitudes (central region maximum = 83; northern region maximum = 1490), and high accuracy was maintained despite differences in the timing of seasonal peaks. These results indicate that our estimation framework is robust to substantial regional variation in both scale and seasonality.

### 4.2 From forecasts to conflict mitigation

The practical value of bear forecasting lies not in reducing bear occurrences themselves but in providing actionable information that guides human behavior. Our forecasts cannot directly suppress bears’ use of areas outside forests. They can, however, deliver timely alerts that discourage people from entering or approaching forested habitats during periods of elevated risk. For such warnings to be effective, they must be intuitively interpretable. A standardized indicator—the bear index—achieves this clarity. Defining the historical monthly maximum as 100 enables straightforward messages: “Bear activity this month is expected to reach 80% of the past maximum.” Human risk perception systematically relies on past baselines—via reference dependence and anchoring—so defining the historical monthly maximum as 100 leverages well-established cognitive regularities (Kahneman & Tversky, 1979; Tversky & Kahneman, 1974). The bear index captures temporal risk, but risk is not only temporal but also spatial. We provided broad-scale maps of baseline spatial risk and further demonstrated that bear sightings increase the probability of subsequent encounters within a 500 m radius over the following 3 days. Together, these temporal and spatial components provide explicit information on when and where encounters are most likely. This enables efficient management without imposing unnecessary restrictions on human activity. Conventional mechanistic models rely on causal variables. In contrast, temporal dynamics alone can generate practical information for conflict mitigation. Critically, this approach remains operational even when environmental predictors are unavailable—a key advantage where data collection is difficult or costly.

### 4.3 Forecast horizon

While our short-term forecasts demonstrated high practical utility, wildlife management agencies often inquire whether predictions can be extended further into the future. Could refining our approach enable longer-term forecasting? The answer is almost certainly no. Consider a hypothetical “perfect” model that completely explains individual bear behavior—one that fully captures habitat selection and foraging dynamics. Even such a complete model could not forecast bear behavior over extended periods. This fundamental constraint, known as the forecast limit or forecast horizon (Beckage et al., 2011; Petchey et al., 2015; Wesselkamp et al., 2025), explains why perfect models inevitably yield imperfect predictions.

Weather forecasting illustrates this principle well. Despite precise modeling of atmospheric dynamics, predictable periods extend only to about ten days (Krishnamurthy, 2019). The barrier lies not in model inadequacy but in sensitivity to initial conditions (Dietze, 2017). Minute errors in the initial state amplify exponentially; extending forecasts by mere days would require order-of-magnitude reductions in measurement error (Krishnamurthy, 2019). Meteorological systems exhibit error-doubling times of roughly two days, meaning a 1°C initial error can expand to 16°C by day eight. Even if bear behavior could be described perfectly by a model, uncertainty in the system’s initial state remains irreducible. Consequently, long-term bear forecasting faces the same fundamental limits as weather prediction.

Furthermore, our approach explicitly relies on temporal autocorrelation, using recent sighting rates to project near-future activity. Once seasonal conditions shift and the correlation with recent history weakens, forecast skill inevitably declines over longer horizons. In practice, this means our method is well-suited for short-term prediction but inherently ill-suited for long-term forecasting, regardless of further parameter tuning. However, within this short forecast horizon, our temporal ensemble appears to efficiently capture a significant portion of the available predictability, achieving this with a framework that is considerably simpler and less data-demanding than fully mechanistic models.

### 4.4 Complementarity of timescales

Long-term forecasts do exist. Climate-driven ecosystem predictions and extended weather projections fall into this category. However, these methods do not sequentially compute the future from current conditions. Rather, they are macro-scale models predicated on the assumption that response variables change gradually in response to slowly varying explanatory variables (Beckage et al., 2011). While day-by-day calculations cannot accurately predict the temperature one year ahead, making probability statements—such as whether next year’s maximum temperature will exceed the historical average—remains feasible by utilizing slowly varying drivers like El Niño or atmospheric CO concentrations (Krishnamurthy, 2019; Shen et al., 2023).

Yet, within ecological forecasting frameworks, discussion of what can and cannot be predicted remains scarce (Beckage et al., 2011; Oliver & Roy, 2015). Clearly recognizing these limits prevents the offering of fundamentally impossible predictions and thereby strengthens scientific credibility (Wesselkamp et al., 2025). COVID-19 forecasting clearly demonstrated this principle: as multiple research groups competed in sequential predictions, both the accuracy and the inherent limits of such models were revealed. Ecological forecasting must likewise acknowledge the boundaries of predictability and uncertainty while still providing information that is useful for management. Our short-term forecasting approach and conventional mechanism-based seasonal predictions operate on different timescales and principles, and thus play complementary roles. Short-term forecasts cannot capture long-term trends, whereas seasonal or longer-term predictions inevitably miss short-term fluctuations. The coexistence of both approaches is essential for establishing robust predictive systems.

## ACKNOWLEDGEMENTS

We thank Kaori Muramatsu for her invaluable assistance with data preparation and formatting.

## AUTHOR CONTRIBUTIONS

Takeshi Honda: Conceptualization, methodology, data curation, formal analysis, investigation, visualization, writing-original draft, writing-review and editing. Chizuru Kozakai: Conceptualization, writing-review and editing.

## STATEMENT ON INCLUSION

Our study relies entirely on secondary data produced by prefectural governments and the Ministry of the Environment in Japan, without new local data collection. All authors are based in institutions within the study region and have long-standing collaborations with local wildlife and agricultural authorities, with whom the forecasting results are shared.

## DATA AVAILABILITY STATEMENT

The data that support the findings of this study have been deposited in the Zenodo repository (reserved DOI: https://doi.org/10.5281/zenodo.17850567) and will be made openly available upon publication of this article.

