## appendix 2 for "Know Today, Know Tomorrow: Ensemble Forecasting of Wildlife Sightings from Temporal Dynamics"

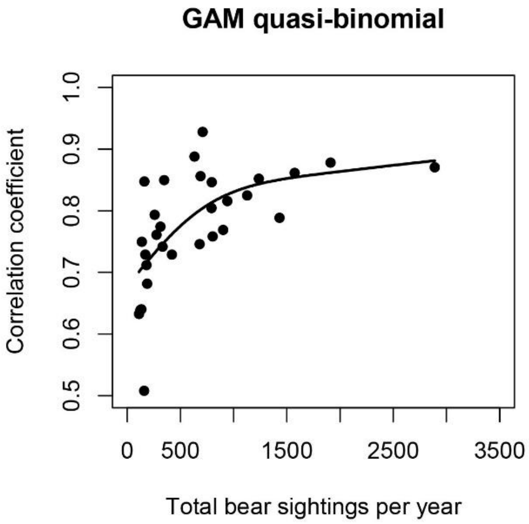


Appendix Figure 1. Relationship between the number of bear sightings and forecast accuracy. In this graph, the forecast represents Ensemble Forecast_base and Forecast_ratio, corresponding to the initial forecast at the beginning of each month. The correlation coefficient, used as an index of forecast accuracy, was treated as the response variable, while the annual number of sightings in each prefecture served as the explanatory variable. The relationship was estimated using a generalized additive model (GAM) with a quasi-binomial distribution.

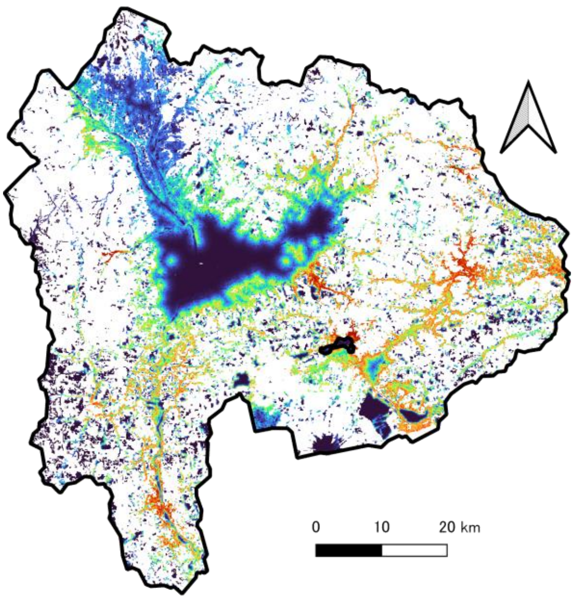


Appendix Figure 2. Spatially estimated baseline risk of bear sightings. This map illustrates the fundamental spatial distribution of baseline risk, which was used as a reference for modeling short-term consecutive sightings. Red areas indicate higher risk and blue areas lower risk, while white areas represent forests excluded from risk evaluation. With the cutoff tuned to equalize sensitivity and specificity, the resulting classification accuracy was 82.1%.
